# Revisiting resonance-excitation collision-induced dissociation for data-independent acquisition

**DOI:** 10.64898/2026.05.31.729078

**Authors:** Chris Hsu, Lilian R. Heil, Bo Wen, Graeme McAlister, Gennifer Merrihew, Philip M. Remes, Deanna L. Plubell, Rafael Melani, Vlad Zabrouskov, Michael J. MacCoss

**Affiliations:** Department of Genome Sciences, University of Washington, 3720 15th Street NE, Seattle, Washington, 98195, United States; Thermo Fisher Scientific, 355 River Oaks Parkway, San Jose, California 95134, United States

## Abstract

Data-independent acquisition (DIA) proteomics relies almost exclusively on beam-type collision-induced dissociation (HCD) because its short activation time supports fast acquisition rates required for DIA. However, HCD requires charge state calibration and is therefore imperfect for mixed-charge DIA isolation windows. Resonance-excitation collision-induced dissociation (reCID) offers a promising alternative to HCD for DIA because the ion activation is effectively independent of charge state. Historically, reCID’s longer activation time has been considered too slow for DIA. Here, we revisit reCID for DIA proteomics using a modified Orbitrap Tribrid Apex MultiOmics mass spectrometer (Apex) that recovers acquisition-matrix overhead as additional ion injection time, enabling reCID acquisition rates comparable to HCD. Using matched acquisition rate settings of tryptic HeLa digests, reCID achieved precursor and protein detections similar to HCD when used with Carafe fine-tuned, fragmentation-matched spectral libraries. Library fine-tuning improved reCID precursor detections more than HCD detections, 24% versus 5%, indicating that HCD-trained prediction models are suboptimal for reCID spectra. ReCID also maintained peptide-level quantitative performance, including ions measured per peptide, precision, and accuracy. Across seven NCI cancer cell lines and pooled mixtures, protein abundance rankings were highly conserved between the Apex reCID and HCD methods, and also across Orbitrap Astral Zoom platforms. These results support reCID as a practical fragmentation mode for DIA proteomics.

## Introduction

Tandem mass spectrometry (MS/MS) has been the foundation of bottom-up proteomics for over three decades.^1^ Early peptide sequencing was performed on tandem-in-space instruments, initially four-sector magnetic sector and triple quadrupole mass spectrometers, and later on Q-TOF and Q-FT-ICR hybrids. In these instruments, precursor ions selected in the first mass analyzer are accelerated into a collision cell, and the resulting fragment ions are mass-analyzed in a subsequent stage.^2-5^ In this beam-type collision-induced dissociation (CID), internal energy is deposited into the precursor in a single pass through the collision cell, and the extent of fragmentation is set by the user-defined collision energy. A persistent challenge of beam-type CID is that the collision energy must be high enough to fragment the precursor backbone yet low enough to avoid secondary fragmentation of the resulting product ions, and the optimal value depends on the mass and charge state of the precursor. In practice, the collision voltage offset is selected from a roughly linear relationship with precursor *m/z* and charge, so that any single setting is well matched only to precursors of similar mass and charge.^6,7^

The introduction of three-dimensional and linear ion trap mass spectrometers brought tandem-in-time MS/MS to proteomics and with it a fundamentally different mode of CID.^8-10^ In resonance-excitation CID (reCID), the trapped precursor is selectively excited at its secular frequency, gaining kinetic energy through many low-energy collisions with a helium bath gas over a user-defined activation period. Because product ions have different secular frequencies than the precursor, they shift off resonance as soon as a single backbone bond cleaves and are no longer activated. This narrow-band activation makes reCID a remarkably efficient fragmentation method, with near-complete conversion of precursor to product, while producing clean, information-rich product ion spectra that are largely free of secondary fragmentation.^9,11^

Despite these advantages, reCID has two well-recognized limitations. First, maintaining stable ion trajectories during resonance excitation imposes a low-mass cutoff at approximately one-third of the precursor *m/z*, which excludes the reporter-ion region of isobaric labels such as iTRAQ and TMT and substantially complicates quantitative workflows that rely on those tags.^12,13^ Second, when peptides carry labile post-translational modifications such as phosphorylation or glycosylation, the lowest-energy dissociation pathway is often neutral loss of the modification rather than backbone cleavage, and reCID spectra of modified peptides can be dominated by a single neutral-loss ion that provides little sequence information.^14^ These limitations motivated a return to beam-type fragmentation on hybrid ion trap-Orbitrap instruments, first demonstrated within the C-trap itself^15^ and subsequently optimized in a dedicated ion-routing multipole behind the C-trap, where it was shown to provide the high-quality low-mass reporter ions and intact phosphopeptide backbone fragments needed for quantitative phosphoproteomics.^16^ This implementation, beam-like CID in an ion-routing multipole (aka HCD), has since become the de facto fragmentation method for Orbitrap-based proteomics.

The dominance of HCD has been further reinforced by the rise of data-independent acquisition (DIA), in which broad precursor isolation windows are systematically fragmented to provide unbiased and reproducible coverage of the precursor mass range.^17,18^ As DIA cycle times have continued to shrink, reCID, with its inherently longer activation times, has been considered too slow to keep pace with modern DIA acquisition rates. Yet, HCD is itself imperfectly matched to DIA: a single normalized collision energy is applied to each isolation window, but those windows routinely contain co-isolated precursors spanning multiple charge states. The applied energy is at best optimal for one charge state and yields under- or over-fragmented spectra for the others, degrading the quality of co-isolated species.^19^

Recent advances in ion pipelining developed for the Orbitrap Apex Tribrid MultiOmics mass spectrometer (MS), also referred to as “Apex”, together with improvements in the linear ion trap, now enable reCID acquisition rates that are comparable to HCD. This eliminates the speed disadvantage that had previously precluded reCID’s use in DIA. The result motivates a fresh look at reCID for DIA proteomics, where two of its intrinsic properties are particularly attractive. Because resonance excitation depends on the secular frequency of the precursor rather than the kinetic energy imparted in a single pass through a collision cell, reCID is effectively charge-state invariant and fragments co-isolated precursors of different charges with comparable efficiency. In addition, reCID of tryptic peptides produces a more intense b-ion series than HCD, providing complementary sequence information that can aid both qualitative detection and quantitation.

Realizing the analytical benefits of reCID in a peptide-centric DIA workflow requires accurate prediction of fragment ion intensities, since modern DIA search tools rely heavily on AI-predicted MS/MS spectral libraries to score peptide-spectrum matches.^20-23^ Almost all of these prediction models have been trained on HCD spectra and do not capture the distinctive fragmentation patterns of reCID. We address this by using Carafe^24^ to fine-tune predicted spectral libraries on a small set of reCID measurements, so that predicted fragment abundances reflect the activation method actually used during acquisition. Together, these advances enable a direct comparison of reCID and HCD for DIA proteomics on a single instrument platform, and we use that comparison to evaluate the analytical merits of reCID across acquisition rate, peptide and protein detection, quantitative reproducibility, and depth of coverage.

## Methods

### HeLa cell culture and SILAC-labeling

HeLa S3 cells were cultured in standard DMEM supplemented with 10% fetal bovine serum (FBS). For SILAC labeling, a parallel population of cells was grown in DMEM containing 13C6, 15N2-L-lysine (Lys8) and 13C6, 15N4-L-arginine (Arg10) using reagents from a commercial kit (Thermo Fisher Scientific). Cells were maintained under SILAC-labeling conditions for 10 doublings to ensure >98% labeling efficiency. Unlabeled and SILAC-labeled cells were harvested at approximately 70% confluency. The cell pellets were washed with 1x DPBS and pelleted by centrifugation and stored at -80°C until lysis.

### Sample preparation

Cell pellets from all samples, including SILAC-labeled HeLa and unlabeled HeLa conditions and the National Cancer Institute (NCI) cell line panel (A549, CCRF-CEM, COLO205, H226, H23, RPMI 8226, and T47D), were processed using methods as previously described.^25,26^ The pellets were lysed in 2% SDS buffer and sonicated (Branson probe sonicator, setting 2). Protein concentrations were determined by using a bicinchoninic acid (BCA) assay (Pierce, Thermo Fisher Scientific), and lysates were diluted to 1 µg/µL in 1% SDS. Lysates were reduced with 20 mM dithiothreitol for 15 minutes and alkylated with 40 mM iodoacetamide for 30 minutes in the dark. The samples were then digested using the protein aggregation capture (PAC) method.^27,28^ PAC was performed by incubating cell lysates with MagReSyn Hydroxyl beads (ReSyn Biosciences) for five minutes. The proteins were then aggregated and captured on the paramagnetic beads with the addition of 70% acetonitrile and incubated at room temperature for 20 minutes. The beads were washed three times with 95% acetonitrile and two times with 70% ethanol on a magnetic rack. After removal of residual solvent, proteins were digested on-bead with trypsin in 50 mM ammonium bicarbonate at a 1:33 enzyme-to-protein ratio for 3 hours at 47°C. The resulting peptides were dried down in a SpeedVac centrifuge and stored at -80°C until data acquisition. Prior to liquid chromatography and mass spectrometry, the samples were reconstituted in 0.1% formic acid. For the matrix-matched calibration experiments, unlabeled HeLa samples were combined with the SILAC-labeled HeLa peptides to generate the following dilutions: 0, 1, 5, 10, 30, 50, 70, and 100% unlabeled content.

### Evaluation of acquisition rate

The acquisition rate was evaluated on both the Orbitrap Ascend Tribrid MS (Ascend) and the Apex instruments using DIA. The method consisted of only MS/MS spectra acquired with 8 Th DIA isolation windows across a precursor mass range of *m/z* 400-1000, continuously for 0.15 minutes. The Orbitrap resolving power was varied (3250, 5400, 7500, 15,000, and 30,000) for both HCD and reCID fragmentation. For reCID, activation time was fixed at 5 milliseconds. The maximum injection time was varied from 0.004 to 80 milliseconds. To ensure that each MS/MS spectrum consistently reached the specified maximum injection time, a Thermo Fisher Scientific H-ESI source was operated with the spray voltage set to 0 V, as previously described.^26^ Raw data files were converted to mzML format using MSConvert (ProteoWizard v3.0.25073).^29^ The acquisition rate was calculated as the total number of spectra divided by the total acquisition (retention) time.

### Liquid chromatography and mass spectrometry

Liquid chromatography separation was performed on a Vanquish Neo UHPLC system (Thermo Fisher Scientific) configured in trap-and-elute mode, using a trap column (300 µm x 0.5 cm) and an Optispray PepMap Neo cartridge analytical column (150 µm x 15 cm). The column temperature was maintained at 45°C, and the ion source was operated at 2250 V. The mobile phase solvents consisted of 0.1% formic acid in water (A) and 0.1% formic acid in 80% acetonitrile (B). Peptides were separated using a 60-minute gradient as follows: 2-35% B over 57 minutes, 35-55% B over 0.5 minutes, 55-99% B over 0.5 minutes, and held at 99% B for 2 minutes. Chromatographic separation was performed at 1.3 µL/min, and column washing was performed at 2.0 µL/min.

Data for HCD and reCID fragmentation comparisons were acquired on a modified Apex instrument. The same instrument could be operated in two modes: (1) a standard, commercially available Orbitrap Ascend Tribrid MS, also referred to as “Ascend”, and (2) a prototype configuration with acquisition control behavior equivalent to an Orbitrap Apex Tribrid MultiOmics MS. The Apex configuration incorporates the improved acquisition timing, scan-matrix scheduling, parallel ion injection control, and Orbitrap signal processing capabilities. Unless noted, all comparisons were performed using matched acquisition parameters between the Ascend and Apex configurations.

MS1 spectra were acquired in the Orbitrap analyzer at 60,000 resolving power over an *m/z* range of 395-1005 and standard AGC target. MS/MS spectra were acquired in DIA mode with 8 Th unstaggered or staggered isolation windows, precursor *m/z* range of 400-1000, 15,000 resolving power with a 1000% AGC target, maximum injection time of 45 milliseconds, and fragment ion *m/z* range of 200-2000. DIA isolation window schemes were experiment-specific. Ascend versus Apex comparisons and normalized collision energy (NCE) optimizations were acquired using unstaggered 8 Th isolation windows. Staggered window DIA comparisons between HCD and reCID on the Apex were acquired using 8 Th staggered isolation windows through a tMSn method. Matrix-matched calibration curve (MMCC) experiments, as well as the NCI7 dataset, were acquired using staggered DIA windows.

NCI7 data were also collected on the Orbitrap Astral Zoom mass spectrometer, referred to as “Orbitrap Astral Zoom HCD.” MS1 spectra were acquired in the Orbitrap analyzer at 60,000 resolving power over an *m/z* range of 395-1005 and standard AGC target. MS/MS spectra were acquired in DIA mode with the Astral analyzer using optimized 3 Th DIA isolation windows and a precursor *m/z* range of 400-1000 as described previously.^26^ HCD collision energy was set to 28 NCE, and fragment scan range of *m/z* 200-1960, 200% AGC target, and 6 milliseconds maximum injection time.

### DIA data search and Skyline post-processing

DIA raw files were processed using Carafe (v2.0)^30^ using the GUI interface to generate a fine-tuned library using a representative run (HeLa or NCI7 pooled sample) from the corresponding experiment. Both fragment ion intensity and retention time predictions were fine-tuned. HCD and reCID datasets were fine-tuned separately so spectral libraries reflected the fragmentation behavior of the method that was used during data acquisition. DIA searches were performed using DIA-NN (v2.3.2) through the Carafe workflow or DIA-NN graphical user interface. For the comparison of library strategies, each dataset was analyzed using both the DIA-NN library-free workflow, in which a predicted spectral library was generated directly from a human FASTA database with DIA-NN’s built-in model, and a Carafe fine-tuned spectral library workflow. Unless otherwise noted, downstream comparisons were performed using the Carafe fine-tuned libraries. DIA-NN search parameters are as shown: trypsin specificity with up to one missed cleavage, fixed carbamidomethylation of cysteines, and precursor charge range of 2+ to 3+. The following DIA-NN options were also enabled: match between runs (MBR), protein inference, unrelated runs, cross-validated neural networks, global normalization, and IM/RT/IDs profiling.

Raw files acquired with staggered DIA isolation windows were converted to mzML and demultiplexed using MSConvert in ProteoWizard (v3.0.25073) with vendor peak-picking and overlap-window demultiplexing using a 10 ppm mass-error tolerance. Non-staggered DIA files acquired on the Ascend, Apex, and Orbitrap Astral Zoom instruments were converted to centroid mzML using vendor peak-picking only. The resulting mzML files were used as input for subsequent DIA-NN searches.

For Skyline data processing, DIA-NN spectral libraries were imported into Skyline-Daily (v26.1.1.097) with the corresponding mzML files. Separate Skyline documents were generated for each experiment and acquisition condition. Unless otherwise noted, MS/MS peak areas were median normalized across runs, and retention time peak boundary imputation was applied in Skyline for missing peak areas.^31^ The matrix-matched calibration curve data was processed in Skyline without median normalization or peak boundary imputation.

## Results

### Modifications to the Orbitrap Ascend Tribrid improve acquisition rate timings

The embedded system of the Orbitrap Apex Tribrid MultiOmics MS instrument was modified to improve ion pipeline and MS matrix execution efficiency by increasing the portion of each acquisition cycle available for ion accumulation. In Tribrid mass spectrometers, ion operations are organized into discrete matrices made up of MS events, times, and device settings. Between these matrices, the MS instrument’s CPUs, FPGAs, and other embedded components must prepare themselves and align with each other prior to executing the next matrix. In previous generation instruments, such as the Orbitrap Ascend Tribrid MS, ion injection could not be parallelized with the time allotted to these embedded system operations. The Apex instrument addresses this limitation, allowing ion injection for the current matrix to extend into the time the embedded systems must consume preparing for the next matrix, thereby converting the unused overhead into additional parallelizable injection time. This scheduling concept is illustrated in Figure 1B for Fourier-transform multistage tandem mass spectrometry (FTMSn), where the top chart in 1B presents the approach on older Tribrids when performing a basic HCD analysis and the middle chart presents the arrangement of ion injection and the embedded system overhead on the new modified instrument. For relatively simple FTMS2 HCD acquisitions, the overhead time associated with preparing the embedded systems for the next matrix is 5.1 milliseconds. However, more complex FTMSn methods, including resonance CID (reCID), require additional ion transfer and manipulation steps because fragmentation occurs in the high pressure trap (HPT) linear ion trap rather than in the HCD cell (Fig 1A). Due to complications arising from the increasingly complex reCID fragmentation matrix, and the need to include Orbitrap ejection and analysis matrix events, it becomes necessary to split out the “Orbitrap analysis stage” into its own matrix (bottom chart of 1B). This change creates an additional period of embedded overhead time in the sequence of events. As a result, even with the modest change of HCD to reCID, the overhead associated with preparing the embedded system increases to 16.0 milliseconds.

**Figure 1.**
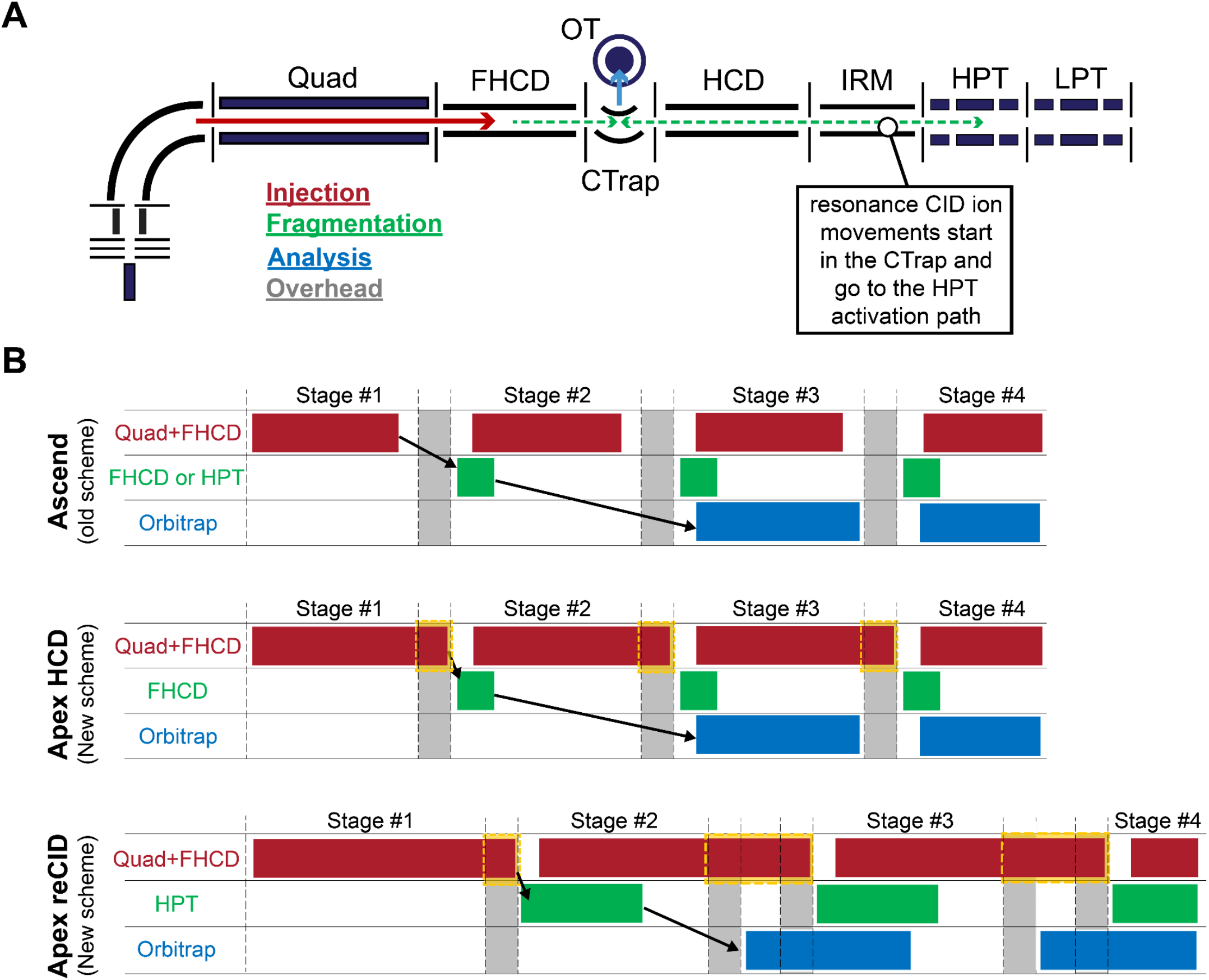
Schematic representation of instrument workflow and matrix sequence for HCD and resonance CID acquisition. **(A)** Instrument schematic showing the ion path through the quadrupole (quad), front-end HCD cell (FHCD), HCD cell (HCD), ion-routing multipole (IRM), high pressure trap (HPT), low pressure trap (LPT), and Orbitrap detector (OT). The arrows indicate the flow of ions through the instrument and the color coding is associated with the operational categories used in the timing diagrams: injection time, fragmentation, Orbitrap analysis, and acquisition-matrix overhead. Note that these categories are simplified descriptions that distill down multiple activities to a single description (e.g., ion movement and fragmentation are simplified to just “fragmentation”). **(B)** Representative matrix sequences comparing the Ascend scheme with Apex HCD and Apex reCID scheme. Each row shows a separate instrument pipeline, and each matrix and overhead period is separated by vertical dashed lines. Red bars indicate injection, green bars indicate fragmentation, blue bars indicate Orbitrap analysis, and grey regions indicate the time consumed by the embedded system overhead. The yellow dashed boxes mark regions where the Apex instrument recovered additional injection time from the overhead (grey) regions. Black arrows indicate an example progression of an acquisition through injection, fragmentation, and analysis. Note these diagrams are meant to be illustrative. The actual size and placement of these boxes may change depending upon factors like ion injection time and Orbitrap transient length.

Because the amount of usable injection time depends on the duration and arrangement of the scan events, we evaluated the acquisition rate as a function of injection time across multiple Orbitrap resolving powers to determine conditions that were suitable for subsequent fragmentation comparisons (Figure S1). For HCD fragmentation, lower resolving powers supported faster acquisition rates at shorter injection times, as expected from the reduced transient duration. In contrast, reCID reached an acquisition rate ceiling of 22 Hz on both the Ascend and Apex instruments at 15,000 resolving power and below. Higher resolving power, such as 30,000, reduced the attainable acquisition rate to under 15 Hz. Based on these data, 15,000 resolving power was selected for the main comparison because it preserved the maximum practical reCID acquisition rate while maintaining higher mass resolution.

At 15,000 resolving power, we compared the acquisition rate with varying injection times for HCD and reCID fragmentation on the Ascend and Apex instruments (Figure 2A). For both fragmentation modes, the Apex maintained the target acquisition rate over a longer injection time range before the acquisition rate began to decline. At an acquisition rate of approximately 19 Hz, the Apex supported a 45 millisecond maximum injection time for both HCD and reCID. Comparable acquisition rates on the Ascend required shorter maximum injection times of 42 milliseconds for HCD and 29 milliseconds for reCID. This difference was reflected in representative HeLa DIA runs, where the Apex methods showed MS2 injection times extending out to the 45 millisecond limit, while the Ascend reCID method was constrained to shorter injection times (Figure 2B). The larger observed improvement for reCID is consistent with the additional scan-matrix overhead required for ion trap fragmentation relative to the HCD workflow.

**Figure 2.**
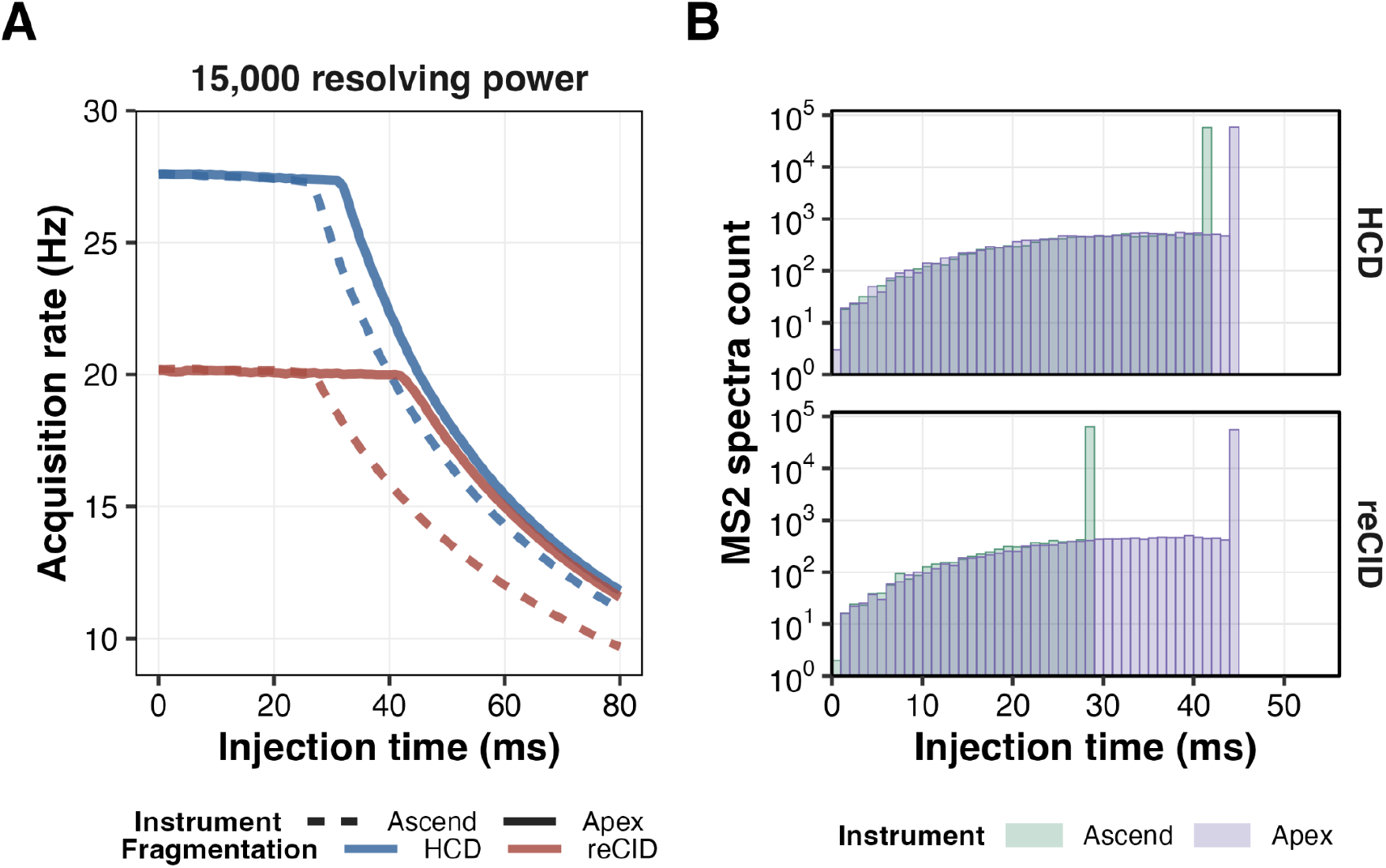
Acquisition rate and injection time improvement of both HCD and reCID fragmentation modes on the Apex and Ascend instruments. **(A)** Acquisition rate comparison between reCID (red) and HCD (blue) on the Ascend (dashed lines) and Apex (solid lines). MS2 spectra were acquired with the spray voltage set to 0 V. **(B)** Distribution of MS2 injection times from a representative run per condition for spectra acquired on the Ascend (green) and Apex (purple) instruments under HCD or reCID fragmentation from a single representative run of HeLa.

Based on these acquisition rate experiments, subsequent comparisons were performed using injection times that produced approximately matched acquisition rates across instruments and fragmentation modes. On the Apex instrument, both HCD and reCID could be operated with 45 milliseconds injection time while maintaining an acquisition rate of approximately 19 Hz. On the Ascend instrument, comparable acquisition rates of 19 Hz required 42 milliseconds for HCD and 29 milliseconds injection time for reCID. These acquisition-rate-matched conditions were used for downstream comparisons so that differences in performance were evaluated at similar acquisition rates and cycle times.

### Resonance CID achieves HCD-like detection depth when fragmentation-matched spectral libraries are used

Before comparing instrument platforms, we optimized fragmentation parameters for HCD and reCID by varying the normalized collision energy (NCE) on the Apex instrument using HeLa tryptic peptides (Figure S2). HCD peptide precursor detections peaked at 28 NCE, whereas reCID peptide precursor detections peaked at 36% NCE. ReCID detection performance was stable from 28 to 44 NCE, while HCD precursor and protein detections declined substantially above 28 NCE (Figure S2A-B). Based on these results, NCE 28 for HCD and NCE 36% for reCID were selected for all downstream instrument and fragmentation comparisons.

Precursor and protein detections were then compared between the Ascend and the Apex using either HCD or reCID fragmentation. Triplicate injections of 500 ng tryptic HeLa peptides were acquired using a 60-minute gradient and 8 Th unstaggered DIA isolation windows. Each dataset was analyzed using two library strategies (see methods): the DIA-NN library-free workflow, in which a spectral library is predicted directly from a supplied FASTA, and the Carafe workflow, in which an experiment-specific library is generated after fine-tuning the retention time and fragment ion intensity prediction models using detections from a representative DIA run for each fragmentation method. The Carafe fine-tuned library or DIA-NN built-in model generated library were then used for subsequent DIA-NN searches.

Carafe fine-tuned libraries increased detections for both fragmentation modes, but the magnitude of improvement was larger for reCID than for HCD (Fig. 3A-B). For the HCD data acquired on the Ascend, precursor detections increased from 81,148 ± 134 with the DIA-NN library-free workflow to 85,669 ± 311 with the Carafe library, corresponding to a 5.6% improvement. On the Apex instrument, HCD precursor detections increased from 91,943 ± 421 to 96,632 ± 491, a 5.1% improvement. Protein detections showed more modest gains for HCD, increasing only 3.0% on the Ascend and 2.8% on the Apex. In contrast, reCID showed larger gains from Carafe fine-tuned libraries. On the Ascend, reCID precursor detections increased from 66,582 ± 175 to 82,584 ± 132, corresponding to a 24.0% increase. On the Apex instrument, reCID precursor detections increased from 79,055 ± 60 to 98,203 ± 79, which was a 24.2% improvement. Protein detections also improved more for reCID than for HCD with an increase of 13.7% on the Ascend and 12.8% on the Apex. These results suggest that reCID fragmentation patterns are less effectively captured by the default DIA-NN library-free prediction workflow, and that fragmentation-matched, fine-tuned library predictions are important for recovering reCID-specific detections. This is consistent with prior results using Carafe.^24^

**Figure 3.**
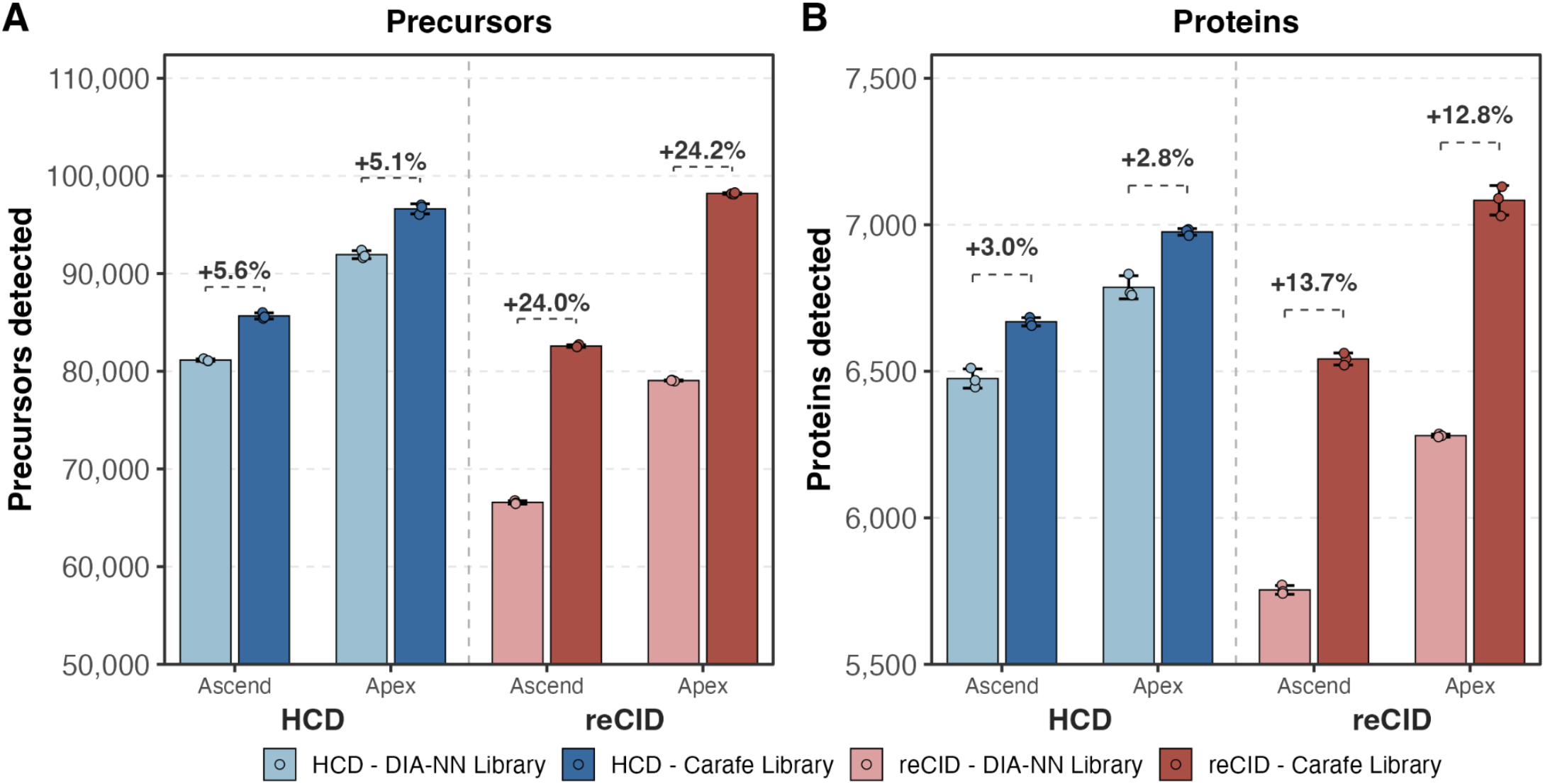
Fine-tuning spectral libraries yields a larger improvement for reCID than for HCD fragmentation. (A) Precursor and (B) protein detections shown for DIA-NN library-free search (light-colored bars) using a spectral library generated from a FASTA compared to Carafe fine-tuned libraries (dark-colored bars), shown for HCD (blue) and reCID (red) on both the Ascend and Apex instruments. Bars represent the mean across replicate injections (n = 3) of tryptic HeLa peptides with error bars indicating the standard deviation and individual replicate values as overlaid points. The bracket annotations indicate the percent increase from using the DIA-NN library-free search to the Carafe library.

When we compare Carafe fine-tuned libraries, the Apex instrument outperformed the Ascend for both fragmentation modes (Figure S3A-C). For HCD, the Apex increased precursor detections by 12.8% and protein detections by 4.6% relative to the Ascend. The improvements were larger for reCID, with precursor and protein detections increasing by 18.9% and 8.3%, respectively. On the Ascend, HCD produced slightly more detections than reCID, whereas reCID on the Apex slightly outperformed HCD. Specifically, the Apex instrument with reCID fragmentation detected 98,203 ± 79 precursors and 7,083 ± 50 proteins, compared to HCD with 96,632 ± 491 precursors and 6,976 ± 11 proteins. This corresponds to a 1.6% increase in precursors and 1.5% increase in proteins.

We examined the overlap in precursor and protein detections across the four acquisition conditions: Ascend HCD, Ascend reCID, Apex HCD, and Apex reCID. The largest intersection of shared detections across all four conditions consisted of 67,531 precursors and 6,562 proteins (Figure S4A-B). This indicates that the majority of detections were conserved across instruments and fragmentation modes. However, the Apex instrument can further extend the detection depth for both HCD and reCID by several thousand precursors and hundreds of proteins. On the Apex instrument, 5,765 precursors and 252 proteins were uniquely detected with HCD, and 7,727 precursors and 299 proteins were uniquely detected with reCID. Rank-abundance analysis showed that these Apex-specific peptide detections were distributed across the low- to mid-abundance range, whereas Apex-specific protein detections were mostly found in the lower-abundance proteins (Figure S5A-B).

Finally, we asked if reCID benefits similarly to HCD from standard DIA acquisition optimizations on the Apex. Using Carafe fine-tuned libraries, we compared 8 Th non-staggered to 8 Th staggered DIA isolation windows, which were demultiplexed prior to DIA-NN analysis to improve precursor isolation specificity.^7,32,33^ DIA staggered windows increased precursor detections from 96,632 to 103,640 for HCD and 98,203 to 105,651 for reCID, corresponding to 7.3% and 7.6% improvements, respectively (Figure S6A). At the protein level, these improvements were 3.4% for HCD and 3.9% for reCID (Figure S6B). These results show that reCID responds to DIA acquisition optimization similarly to HCD. The optimized staggered reCID workflow produced the highest detection depth observed for 500 ng tryptic HeLa peptide injection separated by a 60-minute gradient.

To better understand the MS2-level evidence underlying these detection gains, we evaluated the spectral complexity and the fragment ions selected for quantification. The Apex instrument produced more centroid signals per MS2 spectrum than the Ascend in both HCD and reCID fragmentation modes, indicating that a larger pool of fragment ion signals were available for downstream analysis (Figure S7). Among the fragment ions selected from Carafe fine-tuned libraries, y-ion transition counts slightly favored HCD fragmentation (Figure 4A). However, reCID increased the number of quantitative transitions available for a subset of precursors through b-ion fragments (Figure 4B). HCD showed only a slight benefit in y-ion profiles (Figure 4C), whereas reCID shifted b-ion evidence toward a broader range of fragment m/z values, mostly higher *m/z* b-ions (Figure 4D). These results suggest that reCID increases usable MS2-level evidence mainly by expanding b-ion coverage, without substantially reducing the y-ion information used for DIA-based protein quantification.

**Figure 4.**
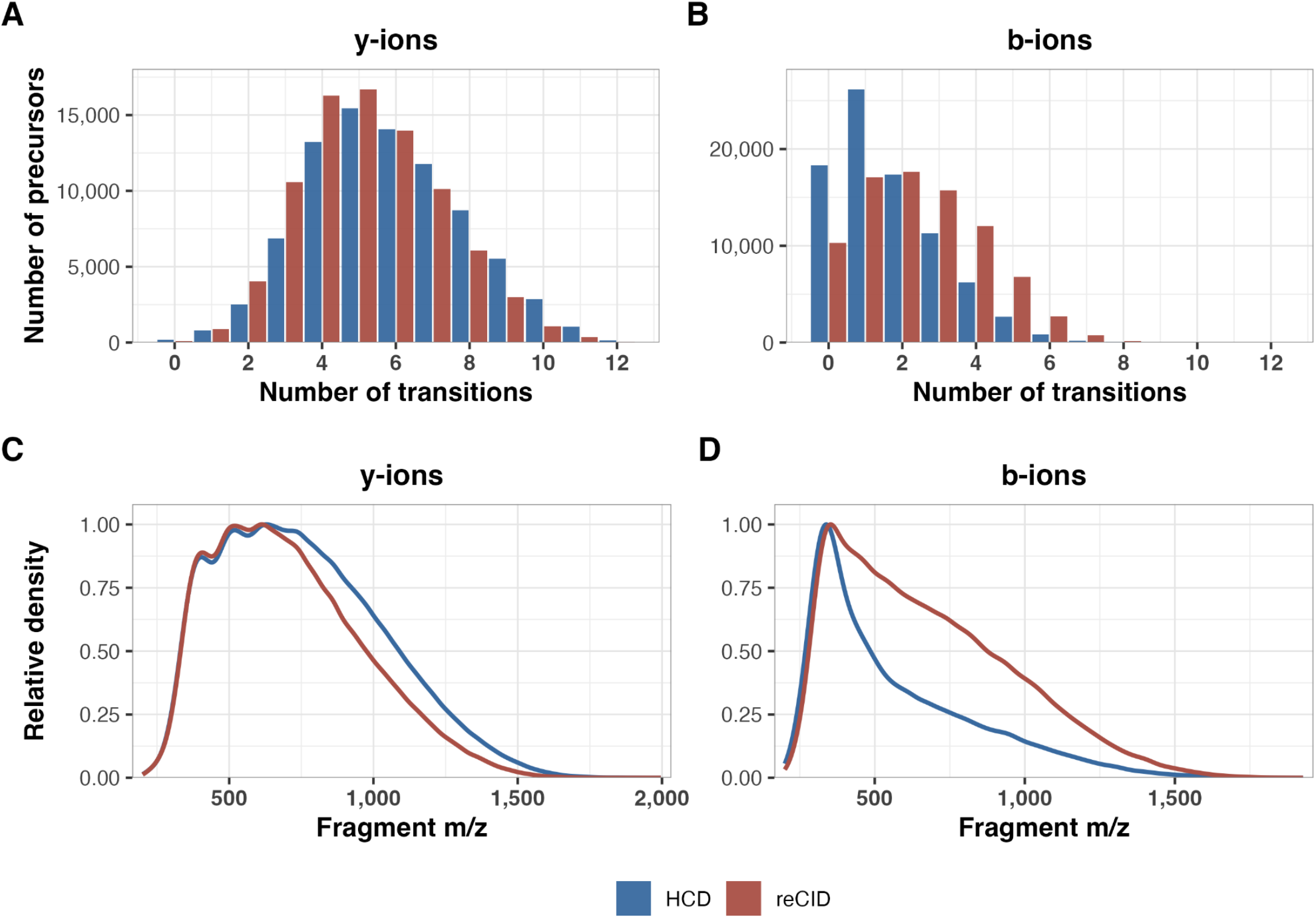
Transition and fragment ion *m/z* differences between HCD and reCID for y- and b-ions on the Apex instrument. Histograms show the distribution of the number of transitions per precursor across shared precursors from the top 12 transitions extracted from Skyline in the Apex instrument using either HCD (blue) or reCID (red) fragmentation. Transition counts are shown for **(A)** y-ions and **(B)** b-ions. Distributions were generated from a representative (n = 1) run for each fragmentation mode using transitions with non-zero peak areas. Relative density distributions of detected **(C)** y-ion and **(D)** b-ion fragment m/z values for shared precursors in a representative replicate on the Apex with either HCD or reCID fragmentation. Fragment ions were only included if they had positive peak areas.

### Assessment of quantitative metrics between reCID and HCD

Having established that reCID can achieve precursor and protein detections comparable to HCD on the Apex instrument, we evaluated whether reCID preserved peptide-level quantitative performance relative to HCD. To enable direct comparison of ion populations across instruments and detectors, raw MS2 signal intensities were converted to ion counts using a calibration constant derived from Glu[1]-Fibrinopeptide B infusion, as previously described.^26^ The Orbitrap MS2 calibration constant was 9.3 for the Ascend and 9.9 for the Apex (Figure S8). These results indicate a similar response between the two instrument modes and demonstrate consistency with previously reported Orbitrap detector calibration factors.

We examined the calibration-corrected apex MS2 total ion counts on both the Ascend and the Apex instrument. On the Ascend, the apex ion counts for both HCD and reCID fell below the corrected AGC target of approximately 101,000 ions, which is consistent with acquisition conditions in which ion accumulation was frequently limited by the maximum injection time (Figure S9A). In contrast, apex ion count distributions on the Apex instrument approached the AGC target with a narrower distribution. This improved ion utilization was supported by the MS2 injection time distributions, which showed a greater use of longer injection times (45 milliseconds on the Apex versus 42 ms on Ascend HCD or 29 ms on reCID) (Figure 2B).

To assess whether this improved ion utilization affected quantitative performance, we compared the calibration-corrected distributions of LC peak peptide ion counts (or ions per peptide) for shared peptides between HCD and reCID. The distribution of LC peak peptide ion counts was very similar between both fragmentation modes with median values of 1534 ions for HCD and 1675 ions for reCID (Figure 5A). The number of MS2 points sampled across the chromatographic peaks was also identical between the two fragmentation modes, giving six points across the peak for both HCD and reCID (Figure S9B). Therefore, reCID and HCD produced comparable peptide-level ion counts with the Apex instrument. Consistent with the similar ion counts and points across the peak, the peptide-level reproducibility was also comparable between fragmentation modes. Across technical replicates, the median peptide CV was 8.2% for HCD and 7.2% for reCID (Figure 5B). The overall CV distributions were similar, indicating that reCID did not compromise replicate-level peptide precision measurements relative to HCD.

**Figure 5.**
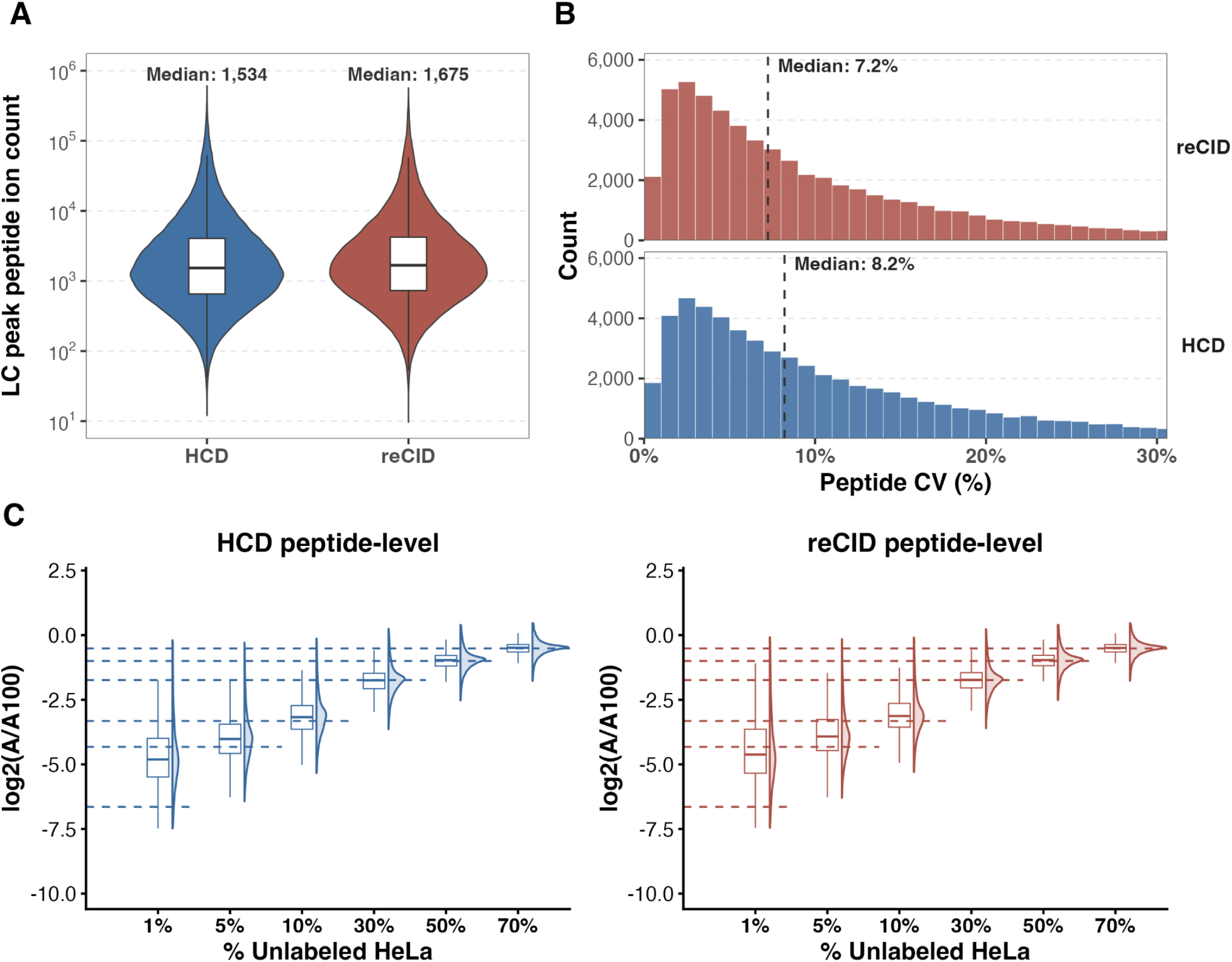
Resonance CID preserves peptide-level quantitative metrics and performance relative to HCD. **(A)** Violin plots with the overlaid boxplots showing distributions of calibration-corrected LC peak peptide ion counts (or ions per peptide) for peptides quantified by both HCD and reCID on the Apex across technical replicates (n=3). Median ion counts are indicated above each distribution. **(B)** Histogram distributions of peptide-level coefficients of variation (CV) across technical replicates for reCID (top, red) and HCD (bottom, blue) on the Apex instrument. Dashed lines and text indicate the median peptide CV for each fragmentation mode. **(C)** Peptide-level quantitative accuracy for an unlabeled HeLa dilution series analyzed by HCD (left, blue) and reCID (right, red). Peptide abundances were normalized to the 100% unlabeled HeLa condition and plotted as a log2(A/A100), where A is the measured peptide abundance at each dilution and A100 is the abundance at 100% unlabeled HeLa. Dashed horizontal lines indicate the expected log2 ratios.

Lastly, we evaluated the quantitative accuracy using a matrix-matched calibration curve by diluting unlabeled HeLa tryptic peptides into SILAC-labeled HeLa background peptides to maintain the same amount of injected material. Samples were prepared at eight concentrations (0%, 1%, 5%, 10%, 30%, 50%, 70%, and 100% unlabeled HeLa), and the data acquired with 8 Th staggered isolation windows on the Apex instrument using either HCD or reCID as the fragmentation mode. The peptide abundances were normalized to the 100% unlabeled HeLa condition. For both HCD and reCID, peptide-level ratios followed the expected dilution trend across the dilution series and gave similar distributions between fragmentation modes (Figure 5C). At the lowest dilution level of 1%, both methods showed broader distributions and ratio compression, which is consistent with previous results on another Orbitrap Tribrid MS.^34^ Taken together, these results show that reCID maintains consistent peptide-level ion counts and quantitative performance that is comparable to HCD on the Apex instrument.

### ReCID captures HCD-based proteomic signatures across NCI samples

To determine if reCID on the Apex preserves biologically meaningful protein abundance measurements beyond benchmarking HeLa samples, we acquired data on seven NCI cancer cell lines and two pooled mixtures, NCI4 and NCI7, using the Apex HCD or Apex reCID in technical triplicate injections of 500 ng per sample by DIA. The NCI4 mixture contained A549, H23, T47D, and CCRF-CEM, and the NCI7 mixture contained those same four cell lines plus COLO205, H226, and RPMI 8226. HCD and reCID acquisitions were performed using 8 Th staggered DIA isolation windows. Across the individual cell lines and pooled mixtures, the Apex reCID produced protein detections similar to the Apex HCD, indicating that proteome coverage was also maintained across diverse cancer cell lines (Figure 6A).

**Figure 6.**
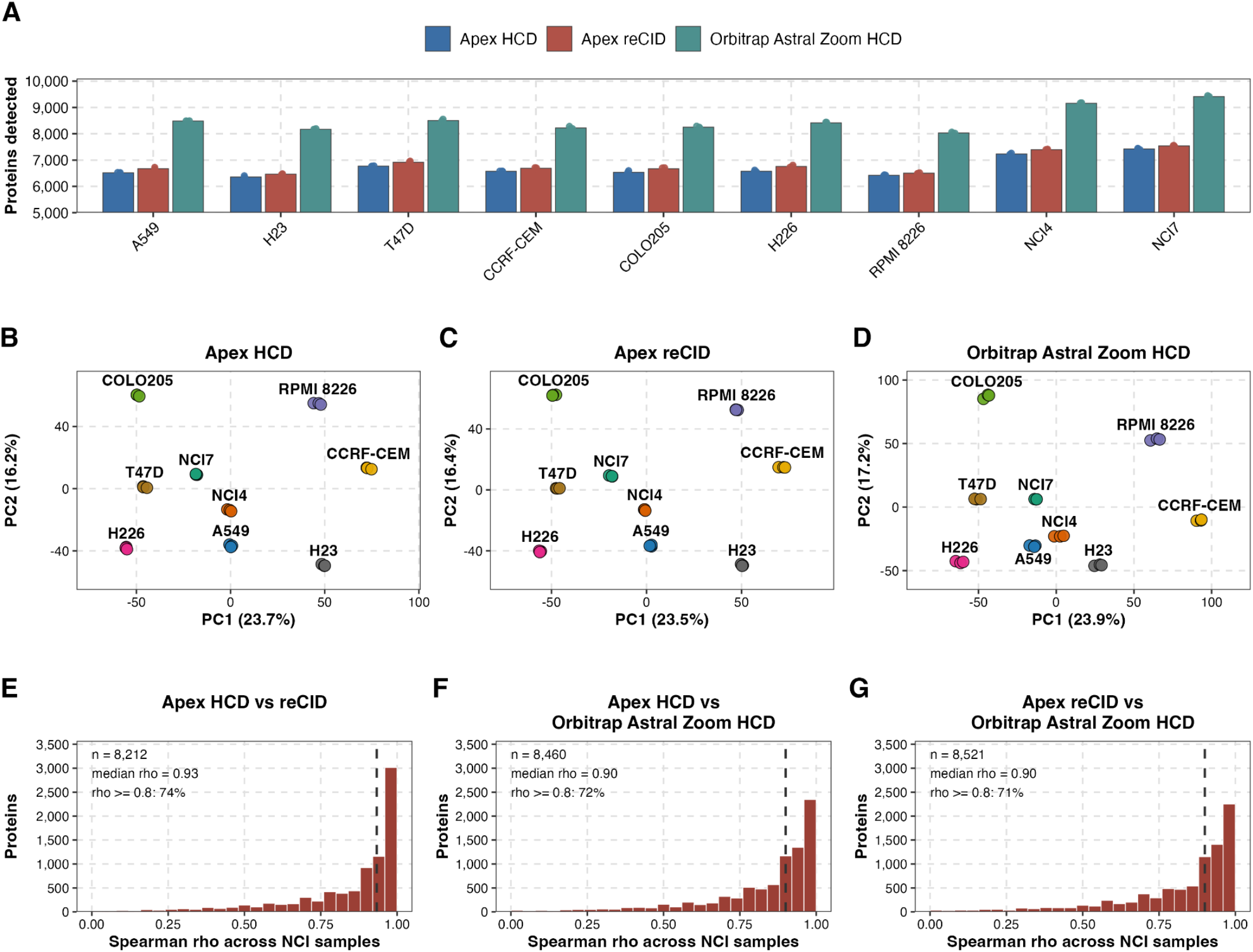
ReCID preserves protein abundance structure across NCI cell-line samples. **(A)** Number of proteins detected in each NCI7 sample for the Apex HCD, Apex reCID, and Orbitrap Astral Zoom HCD. Bars show the mean across replicate injections and the points show individual replicates. **(B-D)** Principal component analysis of protein abundance profiles for Apex HCD, Apex reCID, and Orbitrap Astral Zoom HCD. **(E-G)** Distribution of pairwise Spearman correlations for protein abundance profiles between Apex HCD and Apex reCID, Apex HCD and Orbitrap Astral Zoom HCD, and Apex reCID and Astral Zoom HCD. Correlations were calculated across the nine NCI7 samples using the pairwise intersection of shared proteins for each method comparison.

To assess whether reCID preserved the biological structure of the dataset, we performed principal component analysis (PCA) on the protein abundance measurements from each acquisition method. Apex HCD and reCID showed similar PCA clustering for each of the individual cell lines and pooled mixtures with technical replicates grouping tightly together (Figure 6B-C). These results indicate that reCID retained the same major quantitative relationships among the NCI panel samples observed with HCD. Furthermore, we acquired data on the same samples with the Orbitrap Astral Zoom HCD. As expected, the Orbitrap Astral Zoom HCD detected more proteins across all samples (Figure 6A). Despite the increase in protein depth, the Orbitrap Astral Zoom HCD recovered a similar overall organization of the cancer cell lines and pooled mixtures by PCA, indicating that the biological relationships were consistently observed across different instrument platforms and fragmentation methods (Figure 6D).

To determine whether differences in protein detection depth or acquisition strategy altered the relative quantitative structure of the NCI panel samples, we used pairwise Spearman correlations to test whether each method preserved the rank ordering of the nine samples for each protein. Protein-level rank order was highly preserved between methods, with median Spearman rho values of 0.93 for the Apex HCD versus Apex reCID, 0.90 for Apex HCD versus Orbitrap Astral Zoom HCD, and 0.90 for Apex reCID versus Orbitrap Astral Zoom HCD (Figure 6E-G). Most proteins showed strong rank-order correlation across methods, as 74%, 72%, and 71% of the total proteins, respectively, had a Spearman rho ≥ 0.8. Although the Orbitrap Astral Zoom HCD increased protein detection depth, relative protein abundance patterns across the NCI7 panel remained highly consistent between the Apex reCID and the HCD-based instruments and acquisition methods.

## Discussion

Our results show that reCID can be considered as a practical fragmentation mode for DIA proteomics on the modified Orbitrap Tribrid Apex MultiOmics MS instrument. Historically, reCID has been too slow for DIA because ion trap activation adds timing overhead relative to HCD. The Apex instrument reduces this limitation by improving the pipelining of scan timings for reCID, allowing for more usable parallel ion injection at a matched acquisition rate. As a result, HCD and reCID can now be compared at similar acquisition rates and cycle times. Under these matched conditions, reCID achieved precursor and protein detections comparable to HCD, indicating that the historical speed disadvantage of reCID is no longer a fundamental barrier to its use in DIA.

A major finding is that reCID performance strongly depends on using fragment-matched spectral libraries. Carafe fine-tuning improved detections for both HCD and reCID, but the improvement was much larger for reCID. This is consistent with the expectation that HCD-trained or HCD-like prediction models do not fully capture the fragment ion intensity patterns produced by reCID. Specifically, HCD-trained models underpredict the b-ion intensities and shift the predicted fragment m/z distributions away from what reCID actually produces. More generally, peptide-centric DIA workflows that rely on AI-predicted peptide properties should match the prediction model to the activation method used during acquisition; default HCD-trained models cannot be assumed to transfer to other fragmentation modes.

Importantly, the comparable detection performance of reCID was not obtained at the expense of quantitative performance. On the Apex, reCID and HCD produced similar peptide-level ion counts, CVs, and quantitative accuracy in a dilution series. These results suggest that reCID preserves the core quantitative properties required for DIA proteomics. In addition, the NCI cell line experiments suggest that reCID also maintained protein-level biological structure across diverse samples. PCA clustering and pairwise Spearman correlations demonstrated that protein abundance rank ordering across the NCI panel was highly preserved with median pairwise Spearman rho values of 0.93 for Apex HCD versus Apex reCID, 0.90 for Apex HCD versus Orbitrap Astral Zoom HCD, and 0.90 for Apex reCID versus Orbitrap Astral Zoom HCD.

The potential value of reCID for DIA proteomics lies in its distinct fragmentation physics. Because resonance excitation depends on the precursor’s secular frequency rather than on a charge-dependent collision energy, reCID fragments co-isolated precursors of different charges with comparable efficiency, an inherent advantage for wide DIA isolation windows. In addition, the increased b-ion evidence observed with reCID may provide complementary fragment information for precursor detection and quantification. This added b-ion coverage could be relevant for PTM-centric DIA workflows, where confident localization of phosphorylation, glycosylation, or other modifications depends on fragment ions that distinguish candidate modified residues within the same peptide sequence. Prior comparisons of reCID- and HCD-type fragmentation in phosphoproteomics have shown that fragmentation strategy can substantially affect phosphopeptide and phosphosite datasets,^35^ further supporting the broader idea that alternative fragmentation modes may have PTM-specific advantages. However, this potential should be evaluated directly for reCID, since labile PTMs can also undergo dominant neutral loss under resonance excitation. These features may become especially useful as DIA methods continue to push towards shorter cycle times and more complex spectral prediction search tools or models.

However, there are still important limitations to consider. ReCID retains the low-mass cutoff associated with ion trap resonance excitation. Its utility for labile post-translational modifications, such as phosphorylation or glycosylation, also remains uncertain because the activation can favor neutral loss over backbone fragmentation. Here, we only evaluated reCID for unmodified tryptic peptides using DIA on the Apex instrument. Future work should evaluate generalization to other proteases, post-translational modifications, input amounts, DIA window schemes, and charge-state distributions. More broadly, our results suggest that revisiting alternative fragmentation methods with modern acquisition hardware and fragmentation-aware computational models can expand the set of practical options for DIA proteomics.

## Supporting information

Supplemental Apex reCID

## Data availability

Instrument raw files, database search results, and Skyline files are available on Panorama at https://panoramaweb.org/MacCoss_ModifiedOrbitrapApex.url. R code for statistical analysis and plotting are available on Github (https://github.com/uw-maccosslab/manuscript-dia-recid).

## Acknowledgements

This work was supported in part by National Institutes of Health grants R24 GM141156, U01 DK137097, P30 AG013280, T32 AG066574, and the National Science Foundation Graduate Research Fellowship Program (Grant No. DGE-2140004, B.W.).

## Conflicts of interest

The MacCoss Lab at the University of Washington has a sponsored research agreement with Thermo Fisher Scientific, the manufacturer of the mass spectrometry instrumentation used in this research. Additionally, MJM is a paid consultant for Thermo Fisher Scientific. LRH, GM, PMR, DLP, RM, VZ are employees of Thermo Fisher Scientific.

