## Supplemental Apex reCID for "Revisiting resonance-excitation collision-induced dissociation for data-independent acquisition"

### SUPPLEMENTARY FIGURES:

- **Figure S1.** Acquisition rate across varying injection times and resolving powers.
- **Figure S2.** Optimization of the normalized collision energy (NCE) for HCD and reCID fragmentation.
- **Figure S3.** The Apex instrument improves detections for both HCD and reCID fragmentation.
- **Figure S4.** Overlap of precursors and proteins between reCID and HCD on the Apex and Ascend.
- **Figure S5.** The Apex instrument recovers additional peptides and proteins spanning the lower to mid abundance range compared to the Ascend.
- **Figure S6.** Staggered DIA isolation windows improve precursor and protein detections for both HCD and reCID on the Apex instrument.
- **Figure S7.** MS2 spectral complexity across instruments and fragmentation modes.
- **Figure S8.** Glu[1]-Fibrinopeptide B calibration to ions per second for the Orbitrap detector on Ascend and Apex instruments.
- **Figure S9.** MS2 acquisition metrics for HCD and reCID on the Ascend and Apex instruments.

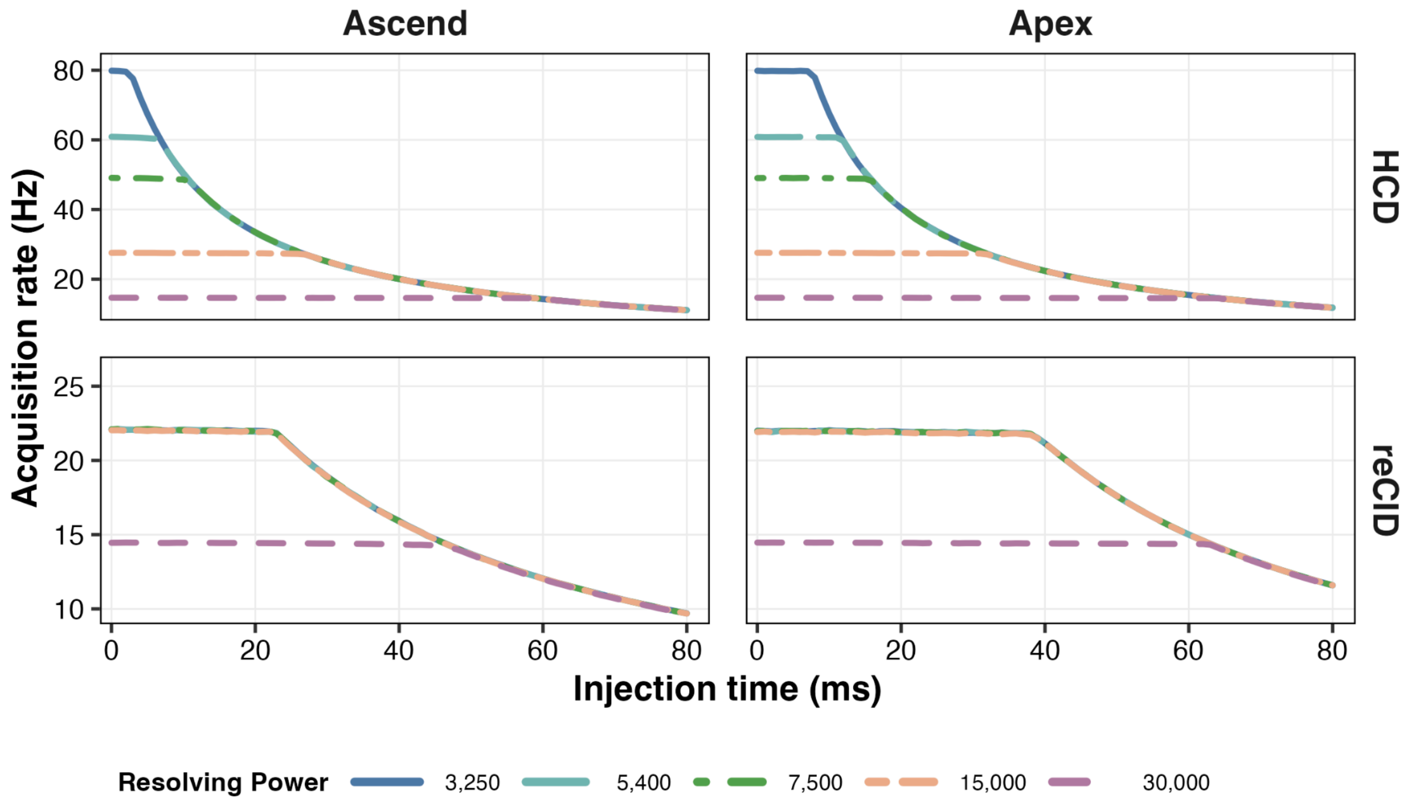


**Supplementary Figure 1. Acquisition rate across varying injection times and resolving powers.** Acquisition rate (Hz) was calculated with injection times from 0 to 80 milliseconds for the Ascend (left) and Apex (right) instruments using HCD (top) and reCID (bottom) fragmentation methods. Columns compare instruments and rows compare the fragmentation pattern. The colored lines represent different resolving power settings of 3250, 5400, 7500, 15,000, and 30,000. Each injection time represents a single replicate measurement.

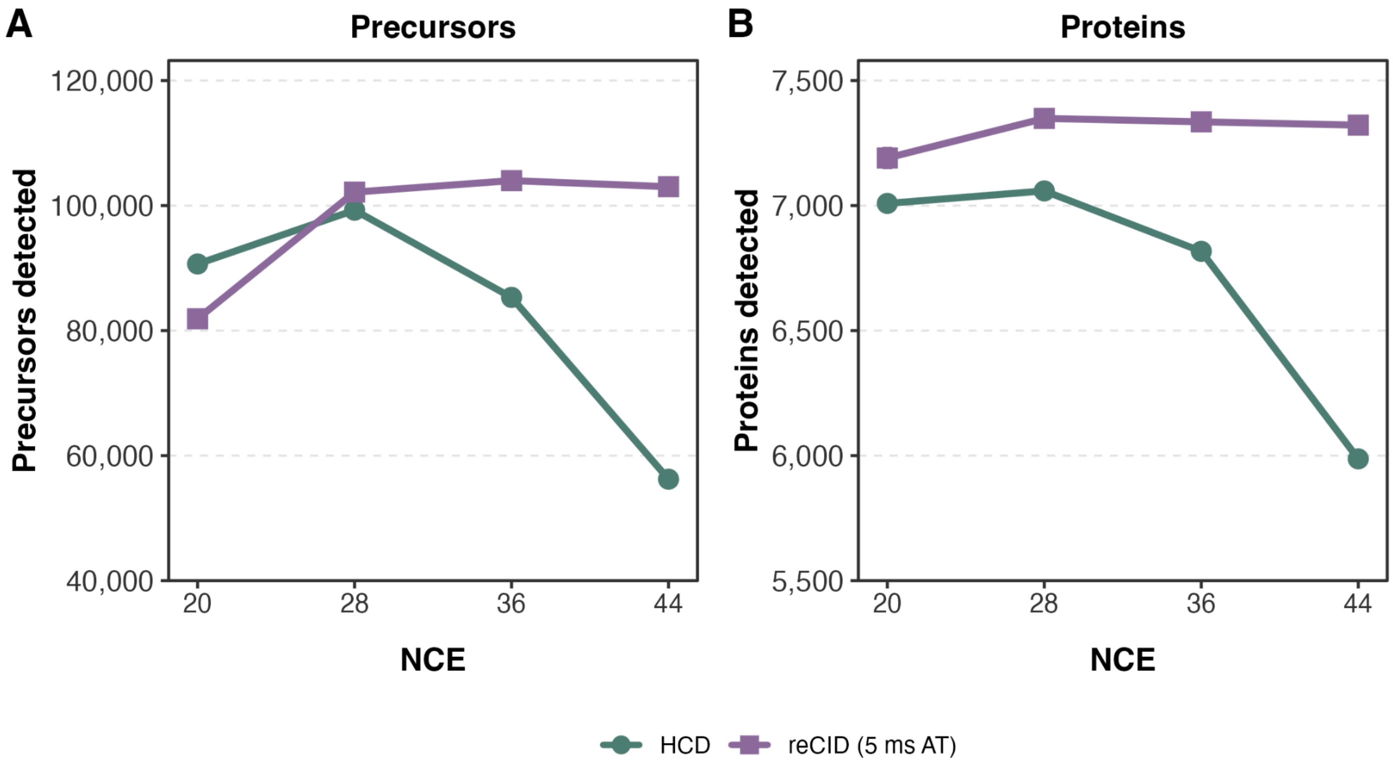


**Supplementary Figure 2. Optimization of the normalized collision energy (NCE) for HCD and reCID fragmentation. (A)** Number of precursors and **(B)** proteins detected as a function of NCE for HCD (green circles) and reCID (purple squares) with 5 milliseconds activation time (ms AT), acquired on the Apex instrument. Detections were retrieved from DIA-NN analysis using a Carafe fine-tuned spectra library. Each point represents the mean of triplicate (n = 3) injections.

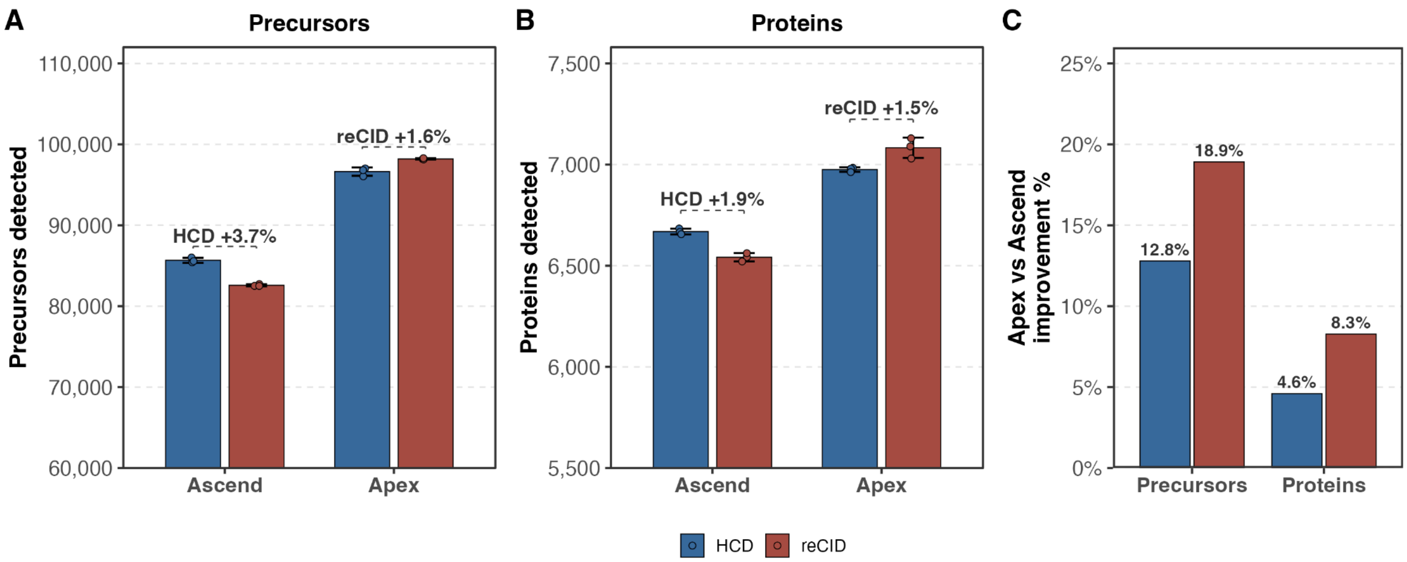


**Supplementary Figure 3. The Apex instrument improves detections for both HCD and reCID fragmentation. (A)** Number of precursors and **(B)** proteins detected using HCD (blue) and reCID (red) on the Orbitrap Ascend or Apex instruments. Bars show the means across triplicate injections of HeLa peptides with error bars represented as the standard deviations and individual replicates overlaid as points. The bracketed annotations report the percent difference within each instrument. **(C)** The percent (%) improvement between the Apex instrument versus the Ascend for either HCD or reCID. Positive percentages indicate the Apex improved more.

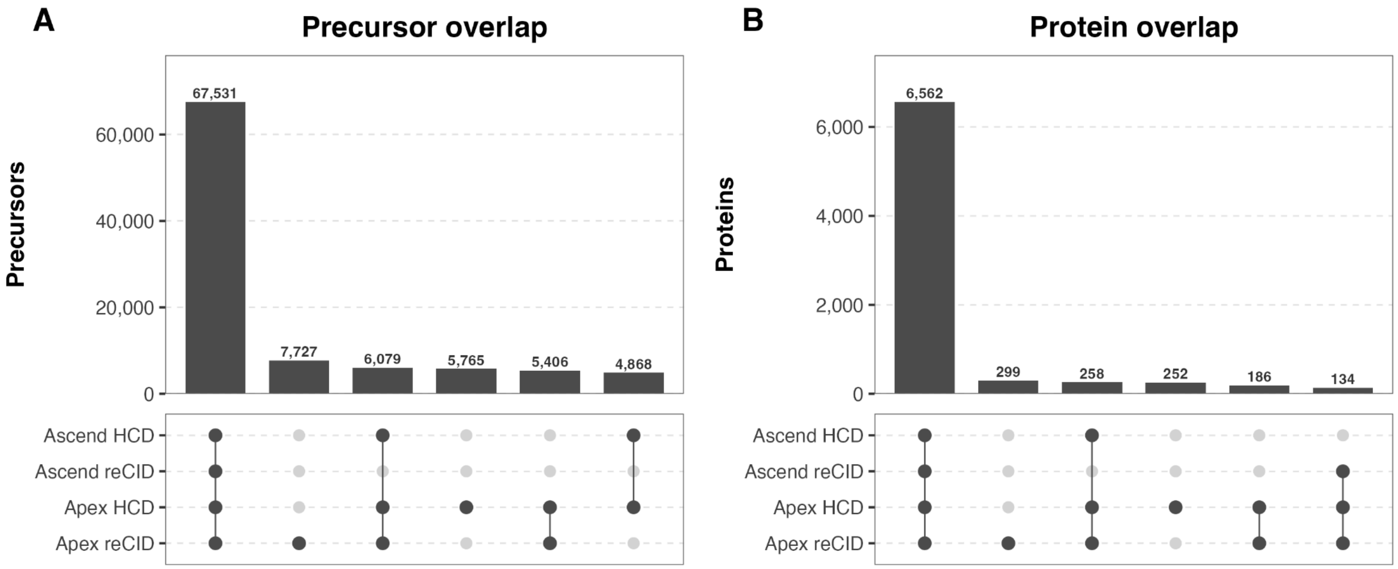


**Supplementary Figure 4. Overlap of precursors and proteins between reCID and HCD on the Apex and Ascend.** UpSet plot showing the overlap of identified (A) precursors and (B) proteins across the four acquisition conditions (Ascend HCD, Ascend reCID, Apex HCD, and Apex reCID) from technical replicates of HeLa tryptic digest (n = 3). The bars indicate the number of precursors or proteins shared among the conditions with the dots indicating the condition below. Only the top six intersections are shown.

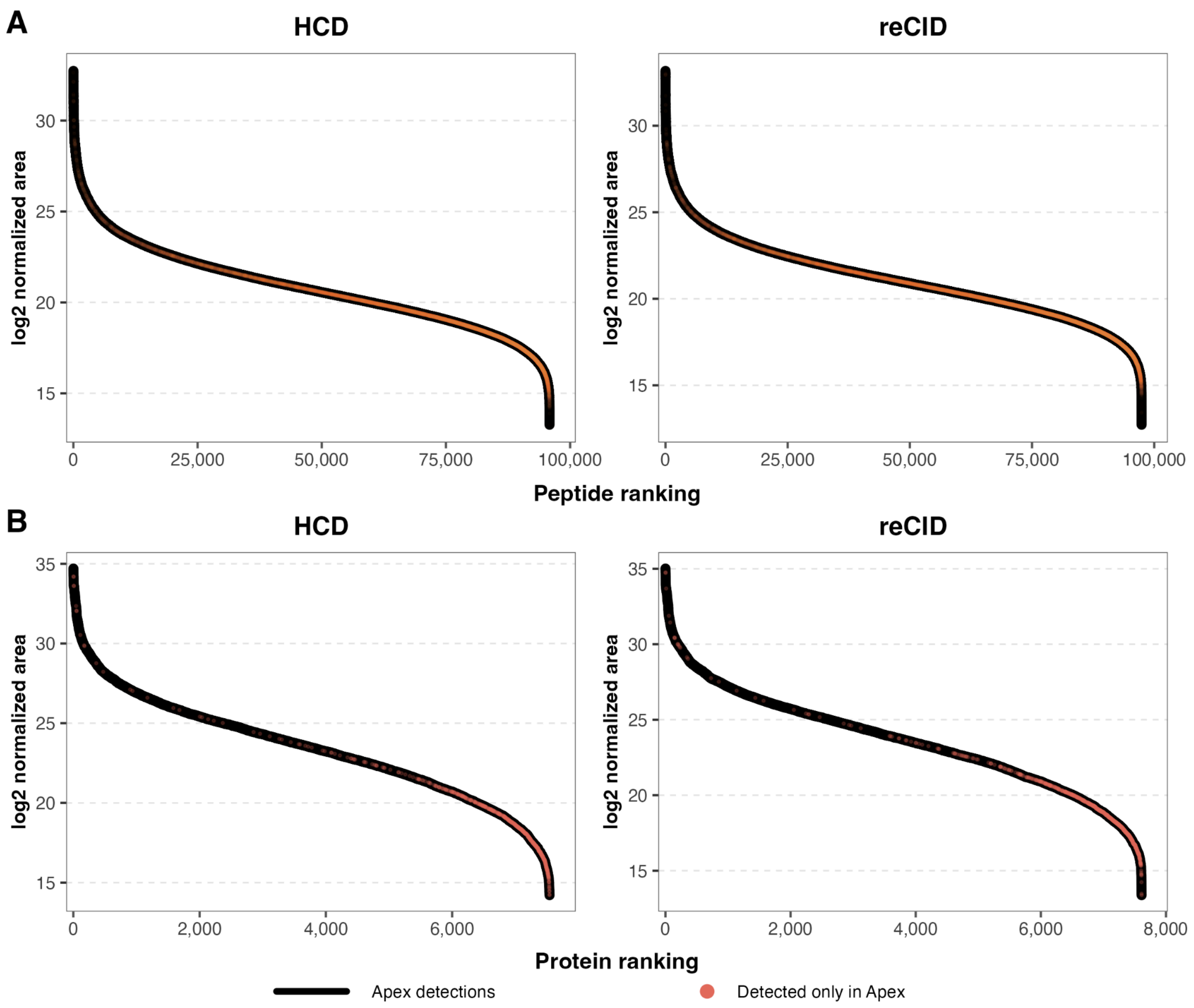


**Supplementary Figure 5. The Apex instrument recovers additional peptides and proteins spanning the lower to mid abundance range compared to the Ascend.** Ranked abundance plots of **(A)** peptides using the log2 mean peak area and **(B)** proteins using the log2 mean protein abundance on the Apex instrument using HCD (left) or reCID (right) from technical replicates of HeLa (n = 3). Peptides or proteins that were exclusively found in the Apex data and not in the Ascend data are noted in red.

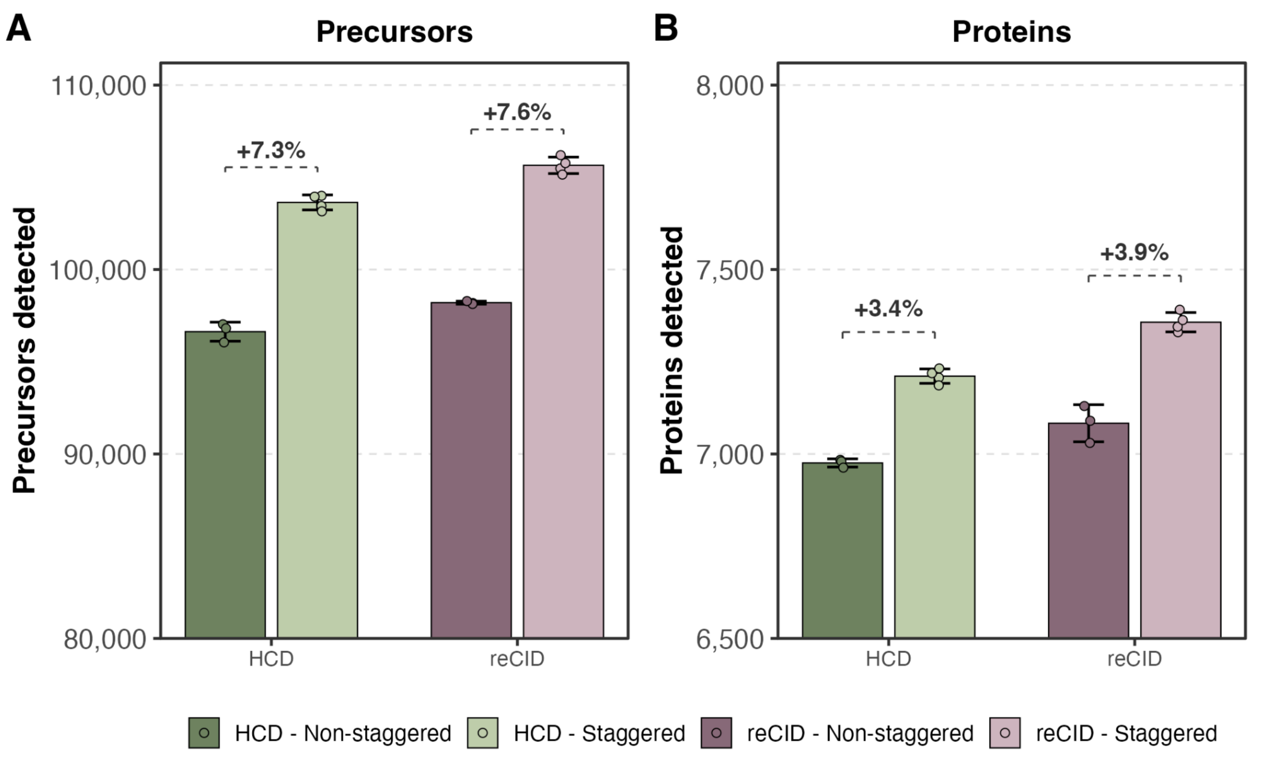


**Supplementary Figure 6. Staggered DIA isolation windows improve precursor and protein detections for both HCD and reCID on the Apex instrument.** Number of **(A)** precursors and **(B)** proteins detected from DIA-NN search using HCD (green) or reCID (purple) fragmentation from technical replicates of HeLa (n = 3). This comparison used 8 Th non-staggered (darker bars) and 8 Th staggered (lighter bars) isolation window schemes. The staggered window analysis was demultiplexed prior to the DIA-NN search. Each bar represents the mean of three technical replicates with individual replicates overlaid as points. The error bars indicate the standard deviation. Percentage values above each pair denote the relative increase in identifications when using staggered versus non-staggered windows within each fragmentation method.

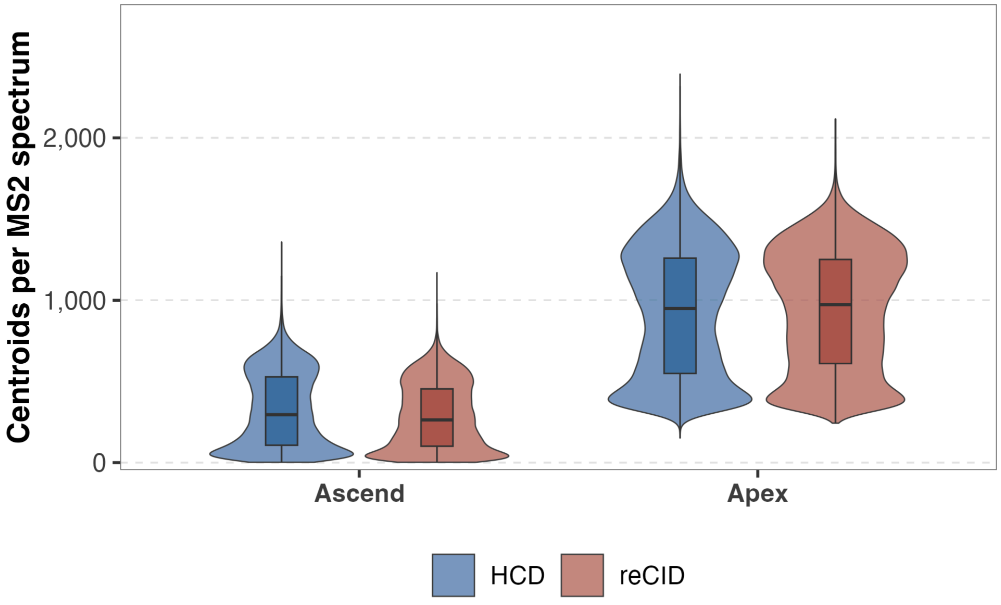


**Supplementary Figure 7. MS2 spectral complexity across instruments and fragmentation modes.** Violin plots with overlaid boxplots show the distribution of centroid signals per MS2 spectrum for HCD (blue) and reCID (red) acquisitions on the Ascend and Apex instruments from one representative run per condition. Each point in the distribution represents one MS2 spectrum.

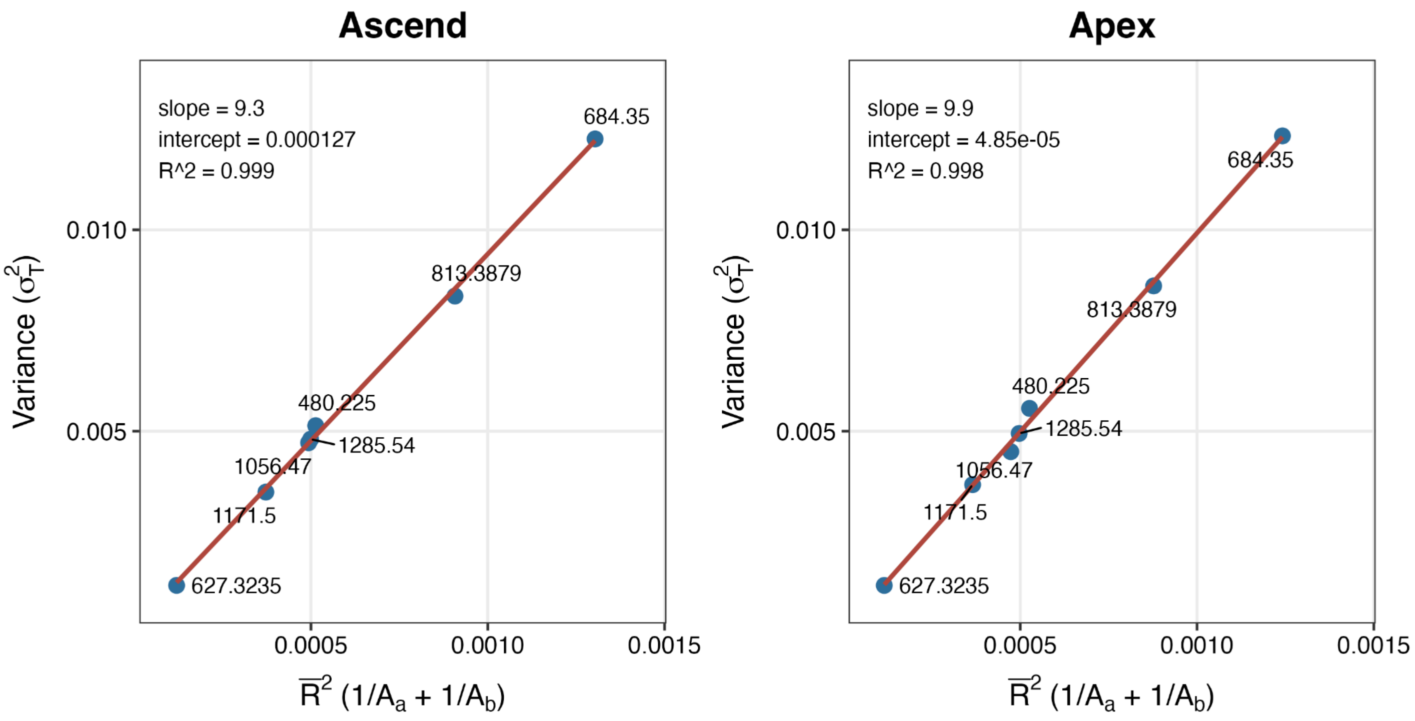


**Supplementary Figure 8. Glu[1]-Fibrinopeptide B calibration to ions per second for the Orbitrap detector on Ascend and Apex instruments.** The Glu[1]-fibrinopeptide B calibration plots showing the squared measurement standard deviation ($\sigma^2_T$) on the y-axis as a function of the mean ratio signal intensity (x-axis) for the Ascend and Apex instruments. The slope represents the α (alpha) factor, the intercept represents other sources of variation, such as detector electronics or signal processing noise, and the $R^2$ values indicate the goodness-of-fit. Each labeled point corresponds to a distinct m/z fragment.

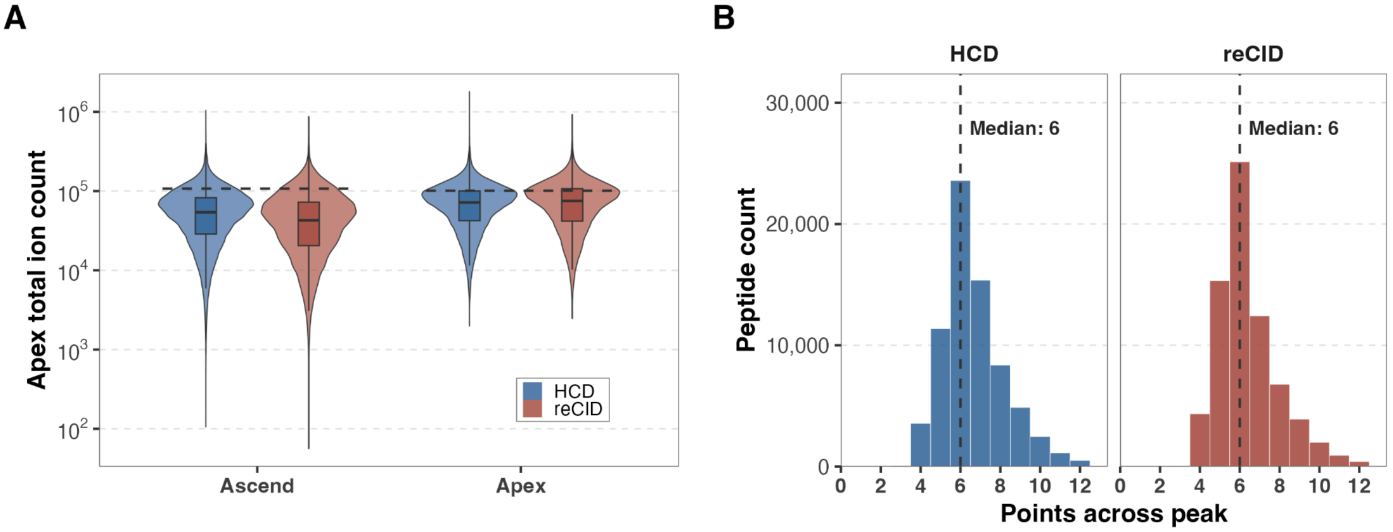


**Supplementary Figure 9. MS2 acquisition metrics for HCD and reCID on the Ascend and Apex instruments. (A)** Violin plots with overlaid boxplots distributions of calibration-corrected apex MS/MS total ion counts for peptides acquired with HCD (blue) or reCID (red) fragmentation on the Ascend or Apex instruments (n = 3, technical replicates). The dashed line indicates the corrected AGC target set by the instrument method. **(B)** Histogram distribution of the average number of points across the chromatographic peak for each shared peptide that were detected by HCD (blue) and reCID (red) on the Apex instrument (n = 3). The dashed line and label indicate the median points across the peaks.
